# Role of sciatic nerve stiffness in surgical decision making and follow up in patients with deep gluteal syndrome

**DOI:** 10.1101/390120

**Authors:** Sava Stajic, Aleksandar Vojvodic, Luis Perez Carro, Jelena Mihailovic, Milos Gasic, Gordana Lukic

## Abstract

The study shows the relevance of sciatic nerve stiffness assessed by strain elastography using ARFI (Acoustic Radiation Force Impulse) for surgical decision making and the follow up of patients with deep gluteal syndrome (DGS). The research focuses on nerve stiffness associated with knee movements in order to determine the degree of nerve entrapment. Neurological examination, MRI of pelvis and electromyography (EMG) were performed as well. The sciatic nerve was scanned by ARFI (strain) elastography during knee movements in patients with DGS (143). In 54 patients surgical treatment was indicated, while 24 of them underwent surgery. The results were based on tissue response to ARFI by color elastogram and stiffness ratio. Diameters of the sciatic nerve in patients with DGS during knee flexion were statistically significantly lower than during extension movement (p<0.01). In patients with DGS (in ones without indication and the ones scheduled for surgery) sciatic nerve stiffness ratio was significantly increased (p<0.01) during knee flexion. Patients scheduled for surgery confirmed increased sciatic nerve stiffness during knee movements, compared with those without indications for surgery (p<0.05). Sciatic nerve recovery after surgery by diameter and stiffness ratio was marked (r=0.881). The correlation between MRI and EMG findings and ARFI nerve stiffness values in patients scheduled for surgery was high (r=0.963). The overall specificity of method was 93.5%, sensitivity was 88.9% with accuracy of 90.6%. ARFI elastography (by strain) is a diagnostic procedure based on nerve stiffness assessment and a useful tool in decision making for surgery and the follow up.

## Introduction

Peripheral nerves tighten and relax during limb movements. Changes in nerve stiffness associated with such movements may be important in understanding various nerve diseases such as deep gluteal syndrome (DGS). The pain originates from the hip joint and the buttock region. Sciatic nerve is positioned underneath the gluteus maximus muscle deep in gluteal region, running between the ischial tuberosity and the greater trochanter of the thigh bone, near the top of the posterior capsule of the hip jont (ball and socket). The nerve may become irritated or inflamed in this tissue. During normal joint movement, the sciatic nerve is flexible enough to stretch and slide with moderate strain or compression. In case of abnormal movements, it may become entrapted or compressed, causing pain. In addition, trauma, injuries or repetitive motions could be causes of pain as well. Sciatic nerve roots emerge from the lumbar plexus within the pelvis and exit through the sciatic notch inferior to the piriformis muscle. Although sciatic nerve is in connection with the superior gluteal and inferior gluteal nerves, the quadratus femoris and the obturator internus, some authors believe that deep gluteal syndrome is a result of sciatic nerve compression caused by piriformis muscle [1,2]. Symptoms vary and may include pain and numbness down the leg [2,3] and they worsen when sitting or standing [3]. Neurological examination, pelvic MRI and electromyography (EMG) are often necessary [4,5]. Acoustic radiation force impulse (ARFI), a low-frequency ultrasound pulse, which is tailored to optimize the momentum transfer to tissue, can be used to create the displacement of tissue [6]. ARFI elastography of sciatic nerve was applied [7–14] with an aim to develop diagnostic possibilities and improve therapeutic modalities. Full testings of sciatic nerve during limb movements are now in the focus of interest. Some authors have noticed sciatic nerve tightening with knee or ankle movements [7,10]. The morphological changes of the sciatic nerve during knee extension movement and knee flexion movement have been observed. The aim of the study is to show the relevance of ARFI elastography (by strain) in nerve stiffness assessment establishing the degree of nerve entrapment in the process of surgical decision making and the follow up.

## Material and methods

The sciatic nerve was scanned by strain elastography using ARFI during knee movements in symptomatic patiens with deep gluteal syndrome (143) over the last three years (from 2015 to 2018). The patients were devided into two groups: group A - without indications for surgical treatment (89) and in group B - scheduled for surgery (54). The recovery of sciatic nerve was followed up in 24 (out of 54) patients who underwent surgery (group C). Monitoring was done three to six months after surgery. The cut off value was calculated in the contol group made of healthy subjects who had been examined (57) and had no signs of DGS (group D). The imaging was performed at the gluteal region. It is important to know that the greater sciatic foramen is an opening in the posterior of human pelvis. It is formed by the sacrotuberous and sacrospinous ligaments and the piriformis muscle passes through the foramen and occupies most of its volume. The greater sciatic foramen is wider in women than in men. The foramen contains seven nerves, with sciatic nerve being the most important one which passes through the infrapiriform foramen. Therefore, the region of interest (ROI) was on the sciatic notch below the piriformis muscle. The sciatic nerve was typically visualized at a depth of 6cm depending on the field of view. By ultrasonography, sciatic nerve was visualized as a hyperechoic, slightly flat oval striped structure. The results by strain elastography using ARFI were presented by color elastograms and relative strain ratio. Nerve excursions were measured during knee extension and flexion test.

The sciatic nerve was scanned using ultrasound elastography equipments (Toshiba Aplio 300). The strain elastography using ARFI was performed by linear probes (5 to 10 MHz). In strain elastography the ARFI pulse replaces the patient or probe movement in order to generate stress on the tissue. This technique may be less user dependent than the manual compression technique. ARFI image can be created by analyzing tissue emplacement. The technique is qualitative (it provides a relative measure of the tissue stiffness in the field of view). The power of ARFI push pulse is limited by guidelines on the amount of energy that can be put into live body, thus limiting the depth of tissue displacement [6]. Methods have been developed that utilize impulsive, harmonic, and steady state radiation force excitations for monitoring the tissue response within the radiation force region of excitation (ROE) and generating images with relative differences in tissue stiffness by acoustic radiation force impulse (ARFI). The tissue displacement resulted in color elastogram, and the calculation of stiffness ratio (SR) was between different regions of interest (ROI).

The obtained data were analyzed using SPSS software (version 22.0 for Windows). The measures of central tendency (arithmetic mean, median), measurements of variability (standard deviation) and relative numbers (structural indicators) were used. The Pearson linear correlation coefficient was used for the analysis of dependence. Statistical hypotheses were tested at the level of statistical significance (alpha level) of 0.05. Finally, ROC analysis was performed.

## Results

Patients with deep gluteal syndrome (DGS), with positive neurological clinical findings (143) were examined by MRI of pelvis (97 out of 143) and by electromyography test (143) (Table I). The patients with DGS were devided according to indications for surgical treatment. In 89 patients surgery was not indicated, whereas 54 were scheduled for surgical treatment. However, 24 patients underwent surgery (Table 1). Regarding the age of patients, there was a statistical significance (p<0.05) between patients without surgical indications and those scheduled for surgery (51.5 to 48.6). Additionaly, the differende in age (p<0.01) between patients who underwent surgery and those who did not (43.5 to 51.5) was found to be statistically significant. The examined patients with DGS were predominatly female (86), while surgical treatment was applied equally to both male and female (Table 1). In surgical decision making MRI findings and electromyography test were decisive (positive in 49 out of 54). Nevertheless, it is important to point out that sciatic nerve ultrasound examination and ARFI elastography test were congruent (Table 1, Table 2 and Table 3).

The results provided by ultrasonography and strain elastography using ARFI were presented by sciatic nerve diameters and by color elastogram and relative stiffness ratio (SR) (Tables 2 and 3). Nerve excursions were measured during knee extension and flexion test. The color imagings were not statistically significant for groups A and B, (patients with and without indication for surgery), due to non-standardized color elastograms and unreliable color images in a certain number of cases. In contrast, the relative differences of sciatic nerve stiffness recorded by ARFI showed results compatible with clinical findings and other diagnostics with exact and practical usefulness. Statistically high diameters of sciatic nerve changed in patients with deep gluteal syndrome during knee movements (Table 2). In knee flexion movements sciatic nerve diameters in group A and B (with and without indication for surgical treatment) were statistically significantly lower (p<0.01) than in extension movements. After surgery, sciatic nerve recovery was markable in follow up (in group C), regarding the nerve diameters in knee extension (p<0.01), while not so high during knee flexion (p<0.05). Significance of nerve stiffness was found during knee movements in groups A and B (patients with and without indications for surgery). In patients with DGS (89), who had been not scheduled for surgery (group A), the relative differences of sciatic nerve stiffness recorded by ARFI was 5.02±1.34SR in knee extension movements, but it was significantly higher (p<0.01) during knee flexion movements 10.27±2.26 SR (Fig 1). What is more, in patients with DGS, scheduled for surgery (group B), the relative differences of sciatic nerve stiffness recorded by ARFI was 6.9±1.34SR in knee extension movements, but significantly higher (p<0.01) during knee flexion movements, 11.8±1.95SR (Fig 2). Statistically significant differences of stiffness ratio was observed (p<0.05) between group A and B (patients not indicated and indicated for surgical treatment). A marked and highly significant decrease of sciatic nerve stiffness ratio during knee movements was detected in patients who had undergone surgery (24). In follow up after surgery (group C) values decreased from 7.32 to 1.32SR during extension, while movements decreased from 11.97 to 4.07SR in flexion (Fig 1 and 2).

**Table 1.** Diagnostics of the subjects with sciatic nerve lesions.

| **Group of Patients** | **Age** | **Sex (male/female)** | **Neurological clinical findings (positive/ examined)** | **MRI (spine and pelvis (nerve lesion/ examined)** | **Electromyography (EMG) (nerve lesion/ examined)** |
| --- | --- | --- | --- | --- | --- |
| A. Patients with deep gluteal syndrome, without surgery (89) | 51.5±8.1 | 35/54 | 89/89 | 18/43 | 45/89 |
| B. Patients with deep gluteal syndrome, scheduled for surgery (54) | 48.6±9.1 | 22/32 | 53/54 | 49/54 | 49/54 |
| C. Patients with deep gluteal syndrome, who underwent surgery (24/54) | 43.5±5.7 | 12/12 | 24/24 | 23/24 | 24/24 |
| D. Healthy subjects (57) | 47.7+-8.6 | 26/31 | 5/57 | 0/5 | 0/9 |

**Table 2.** Diameter of sciatic nerve (mm)

| Group of Patients | Age (y.) | Sex (male/female) | $\bar{x}$ | sd | min | max | p-value |
| --- | --- | --- | --- | --- | --- | --- | --- |
| A. Patients with deep gluteal syndrome, without surgery (89) – knee extension | 51.5±8.1 | 35/54 | 5.96 | 0.69 | 4.3 | 7.3 | <0.01 |
| A. Patients with deep gluteal syndrome, without surgery (89) – knee flexion | 51.5±8.1 | 35/54 | 4.46 | 0.65 | 2.9 | 6.1 |  |
| B. Patients with deep gluteal syndrome, scheduled for surgery (54) – knee extension | 48.6±9.1 | 22/32 | 4.89 | 0.68 | 3.7 | 6.5 | <0.01 |
| B. Patients with deep gluteal syndrome, scheduled for surgery (54) – knee flexion | 48.6±9.1 | 22/32 | 3.98 | 0.68 | 2.6 | 5.4 |  |
| C. Patients with deep gluteal syndrome, who underwent surgery (24/54) - knee extension (*before/after surgery*) | 43.5±5.7 | 12/12 | **4.99/8.05** | **0.70/0.5** | **3.7/7.3** | **6.5/8.9** | <0.01 |
| C. Patients with deep gluteal syndrome, who underwent surgery (24/54) - knee flexion (*before/after surgery*) | 43.5±5.7 | 12/12 | **3.90/4.65** | **0.72/0.24** | **2.6/4.3** | **5.4/5.1** |  |
| D. Healthy subjects (57) – knee extension | 47.7±8.6 | 26/31 | 8.32 | 0.59 | 7.3 | 9.4 | Cut off 7.14 |
| D. Healthy subjects (57) – knee flexion | 47.7±8.6 | 26/31 | 4.79 | 0.31 | 4.4 | 5.6 | Cut off 4.17 |

**Table 3.**
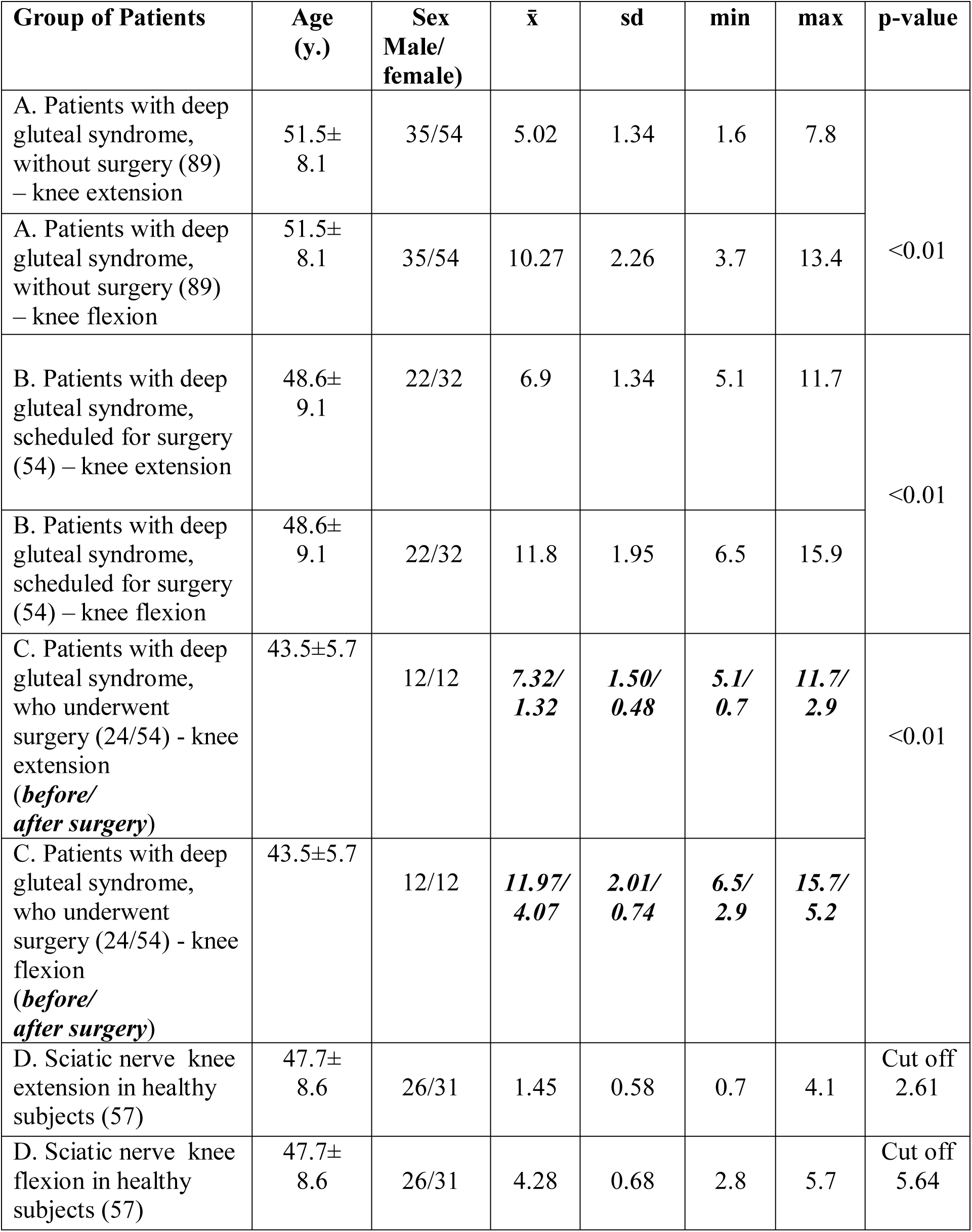
The relative differences of sciatic nerve stiffness by ARFI (SR – Stiffness Ratio)

| Group of Patients | Age (y.) | Sex Male/female) | $\bar{x}$ | sd | min | max | p-value |
| --- | --- | --- | --- | --- | --- | --- | --- |
| A. Patients with deep gluteal syndrome, without surgery (89) – knee extension | 51.5±8.1 | 35/54 | 5.02 | 1.34 | 1.6 | 7.8 | <0.01 |
| A. Patients with deep gluteal syndrome, without surgery (89) – knee flexion | 51.5±8.1 | 35/54 | 10.27 | 2.26 | 3.7 | 13.4 |  |
| B. Patients with deep gluteal syndrome, scheduled for surgery (54) – knee extension | 48.6±9.1 | 22/32 | 6.9 | 1.34 | 5.1 | 11.7 | <0.01 |
| B. Patients with deep gluteal syndrome, scheduled for surgery (54) – knee flexion | 48.6±9.1 | 22/32 | 11.8 | 1.95 | 6.5 | 15.9 |  |
| C. Patients with deep gluteal syndrome, who underwent surgery (24/54) - knee extension (*before/after surgery*) | 43.5±5.7 | 12/12 | *7.32/1.32* | *1.50/0.48* | *5.1/0.7* | *11.7/2.9* | <0.01 |
| C. Patients with deep gluteal syndrome, who underwent surgery (24/54) - knee flexion (*before/after surgery*) | 43.5±5.7 | 12/12 | *11.97/4.07* | *2.01/0.74* | *6.5/2.9* | *15.7/5.2* |  |
| D. Sciatic nerve knee extension in healthy subjects (57) | 47.7±8.6 | 26/31 | 1.45 | 0.58 | 0.7 | 4.1 | Cut off 2.61 |
| D. Sciatic nerve knee flexion in healthy subjects (57) | 47.7±8.6 | 26/31 | 4.28 | 0.68 | 2.8 | 5.7 | Cut off 5.64 |

**Fig 1.**
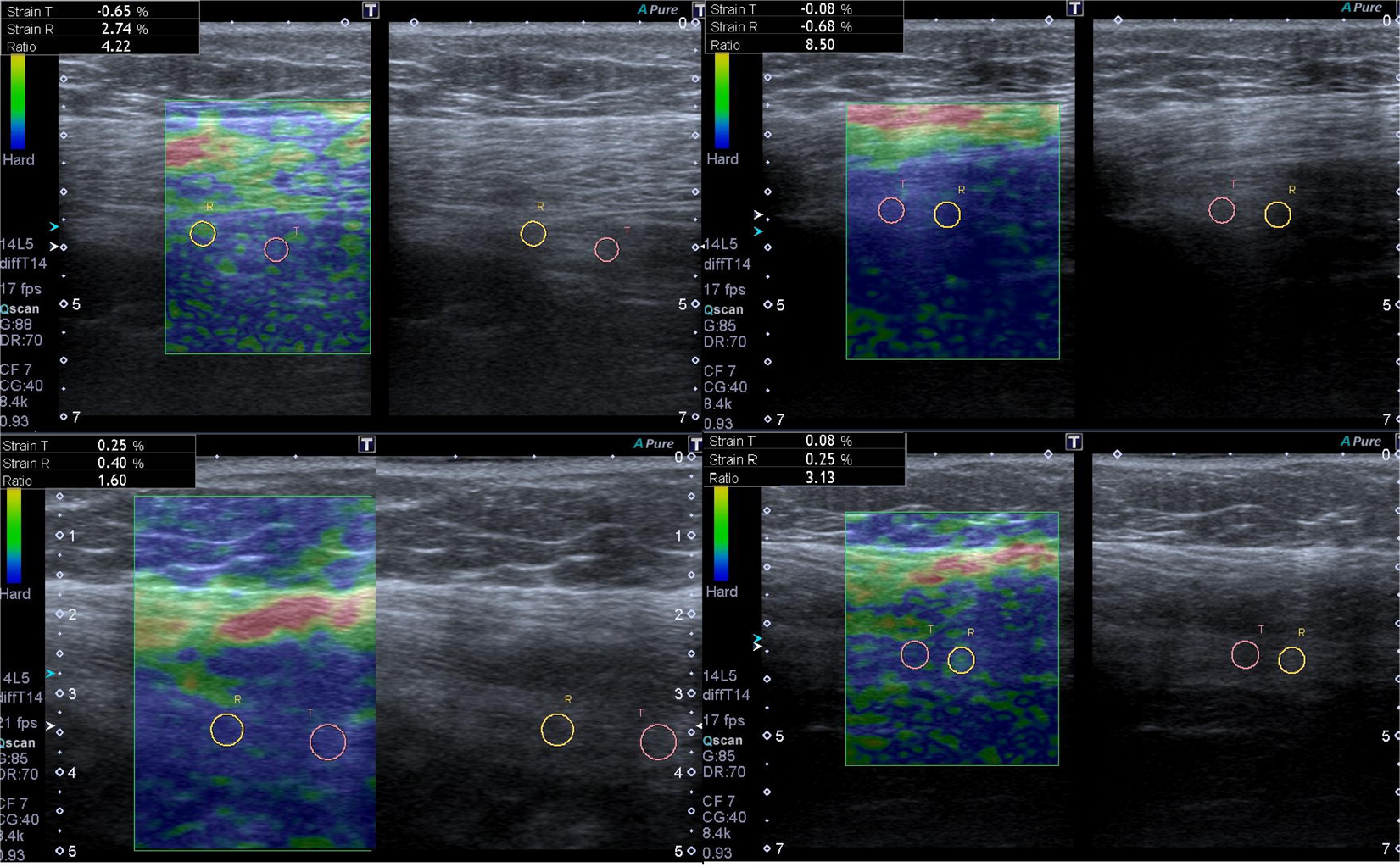
Case 1. The sciatic nerve stiffness before and after surgery.

**Fig 2.**
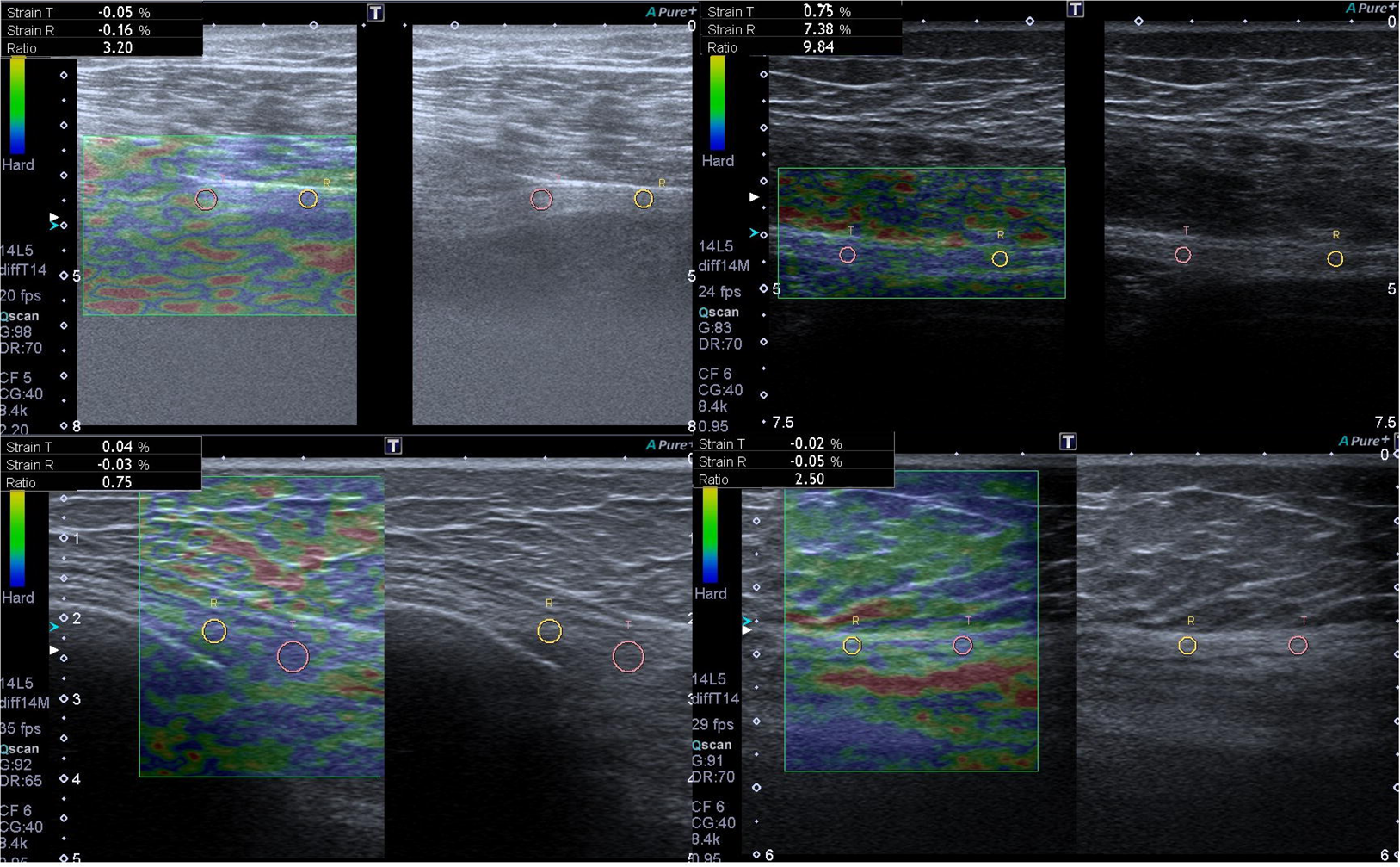
Case 2. The sciatic nerve stiffness before and after surgery.

Case 1 - A 39-year-old female patient, with positive neurological clinical findings, positive MRI of the pelvis, with a compresion of an ovarial cyst on the left sciatic nerve and with moderate nerve lesion found by EMG. The decision was made that, since there were no indications for the surgical release of the sciatic nerve, but she was to undergo a surgery for the ovarial cyst. Relatively high differences of sciatic nerve stiffness were confirmed by ARFI elastography during knee extension and flexion movements (4.22SR and 8.50SR) (above to the left and right). The undefined contour of sciatic nerve, and the differences of sciatic nerve diameters in knee movements have been well-recognized. She underwent a surgery which led to the release of sciatic nerve (1.60SR in extension and 3.13SR in flexion movements) (below to the left and right). Department of Diagnostic Radiology, Clinical Hospital Center „Dr Dragisa Misovic - Dedinje“, Belgrade, 2016.

Case 2 - A 48-year-old male patient with positive neurological clinical findings, positive MRI of lumbal spine and pelvis, with lesion of right sciatic nerve roots and with a severe nerve lesion found by EMG. Relatively high differences of sciatic nerve stiffness were confirmed by ARFI elastography during knee movements (3.20SR in extension, 9.84SR in flexion) before surgery (above to the left and right), with significant decrease after surgery (0.75SR in extension and 2.50 in flexion) (below to the left and right). Department of Diagnostic Radiology, Clinical Hospital Center „Dr Dragisa Misovic - Dedinje“, Belgrade, 2017.

The Pearson linear correlation coefficient was used for the analysis of dependence (Fig 3). The correlation coefficient of sciatic nerve diameter was satisfying in groups A and B (patients without surgery and scheduled for surgery) during knee movements (r=0.557 and r=0.599). In follow up, the correlation coefficient of sciatic nerve diameter was significantly high during knee flexion (r=646) in patients who had undergone surgery (group C). The correlation coefficient of sciatic nerve stiffness ratio during knee movements in patients without surgery (group A) was significant (r=0.683), while in patients indicated for surgery (group B) was moderately significant (r=0.428). The correlation of dependence between diameters and stiffness ratio in group A (without surgery) was significant (r=0.671). The highest significant correlation coefficient was noticed in the analysis of dependence between diameters and stiffness ratio in group C - patients who had undergone surgery (r=0.881). A very high correlation was observed between MRI and EMG findings and ARFI nerve stiffness values in patients scheduled for surgery (r=0.963), as well as in correlation between sciatic nerve diameters and MRI and EMG findings (r=0,833).

**Fig 3.**
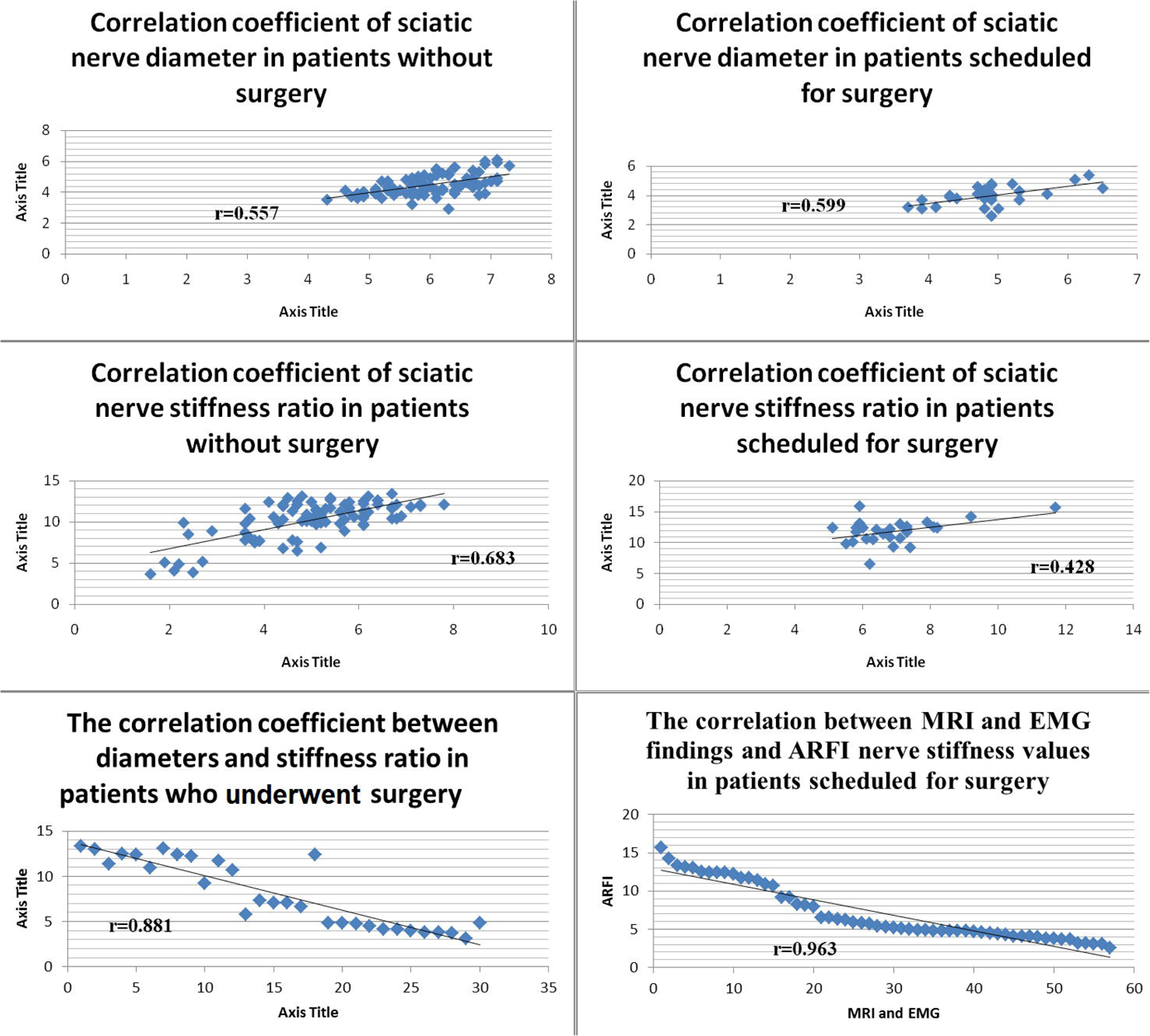
The correlation coefficients.

The Pearson linear correlation coefficient was used in the analysis of dependence between sciatic nerve diameters and stiffness ratio.

In order to calculate ROC analysis it was important to operate by cut off values. In group D made of healthy subjects (57) nerve stiffness during extension movements was 1.45±0.58SR, with cut off 2.61SR, whereas it was 4.28±0.68SR during flexion, with cut off 5.64SR. By ROC analysis (Table 4) overall specificity of ultrasound elastography was 93.5%, in symptomatic patients with DGS, the sensitivity of 88.9% was representative, with high accuracy of 90.6%. [8,9]. The positive predictive value was 82.6%, while the negative predictive value was 96%.

**Table 4.** The operating characteristics by ROC analysis.

|  | % |  | % |
| --- | --- | --- | --- |
| **Specificity of Sciatic nerve during knee extension** | **90.3%** | **Accuracy (%)** | **90.6%** |
| **Specificity of Sciatic nerve during knee flexion** | **96.8%** | **True Positive Values** | **82.6%** |
| **Sensitivity of Sciatic nerve during knee extension and flexion** | **88.9%** | **True Negative Values** | **96%** |

## Discussion

In our study, which investigated patients with DGS, ARFI elastrography test showed high and very useful operating characteristics according to ROC analysis [7,8]. This can be explained by shortening of sciatic nerve diameter, particularly during knee flexion. During such a movement, the nerve was tense, and the stiffness ratio rose markedly.

Ultrasonography detected entrapment and shortnening of sciatic nerve in patients indicated for surgical treatment and the US findings were explicitly correlated with MRI and EMG findings of nerve lesions. It was obviously manifested by diameters of sciatic nerve and by relative differences of sciatic nerve stiffness.

The fibrous bands with nerve entrapment in patients with deep gluteal syndrome significantly decreased the diameter of sciatic nerve in extension, as well as flexion movements, as a result of nerve tightening [11,12]. It was observerd that movements resulted in increased nerve stiffness, particularly prominent during knee flexion movements (Fig 1 and 2). The fact remains that MRI and EMG confirmed nerve injuries. Colour elastograms did not turn out to be particularly helpful, due to lack of their standardization. The obtained images, however, created a possibility for color elastograms to be very useful in some cases, especially in ARFI elastography estimation of the sciatic nerve (Fig 2).

Strain elastography using ARFI showed good performance in follow up of the patients who underwent surgery with regard to the diameter and stiffness ratio. This non-invasive, easily performed and reproducible method could be a very important medical diagnostic tool, especially with respect to cost benefit, having in mind the cost of MRI follow up. Confirmed by ROC analysis as an accurate diagnostic procedure based on the assessment of nerve stiffness, it could be useful in process of surgical decision making and during the follow up.

The prevalence of female population was represented by clinically confirmed cases. There were no significant differences in sciatic nerve diameter with regard to the age. It showed the importance of intraneural stiffness, especially observed in symptomatic patients with morfological changes of the nerves and the surronding fibrous processes (MRI). Therefore, the positioning of ROI was important.

The limits of strain elastography using ARFI depend on the applied techniques, the depth of sciatic nerve and the field of view. The ARFI elastography of sciatic nerve in patients with deep gluteal syndrome was predominantly performed as preoperative decision test (Fig 1 and 2). Upon surgical exploration of the sciatic nerve, a fibrotic tendinous scar beneath the piriformis was found and released. The resection was done from trochanter’s attachment, by separating joint tendon from m. piriformis and m. obturator internus, and by releasing n. peroneus and n. tibialis from fibrous bands and surrounding muscles [15–18] (Fig 4).

**Fig 4.**
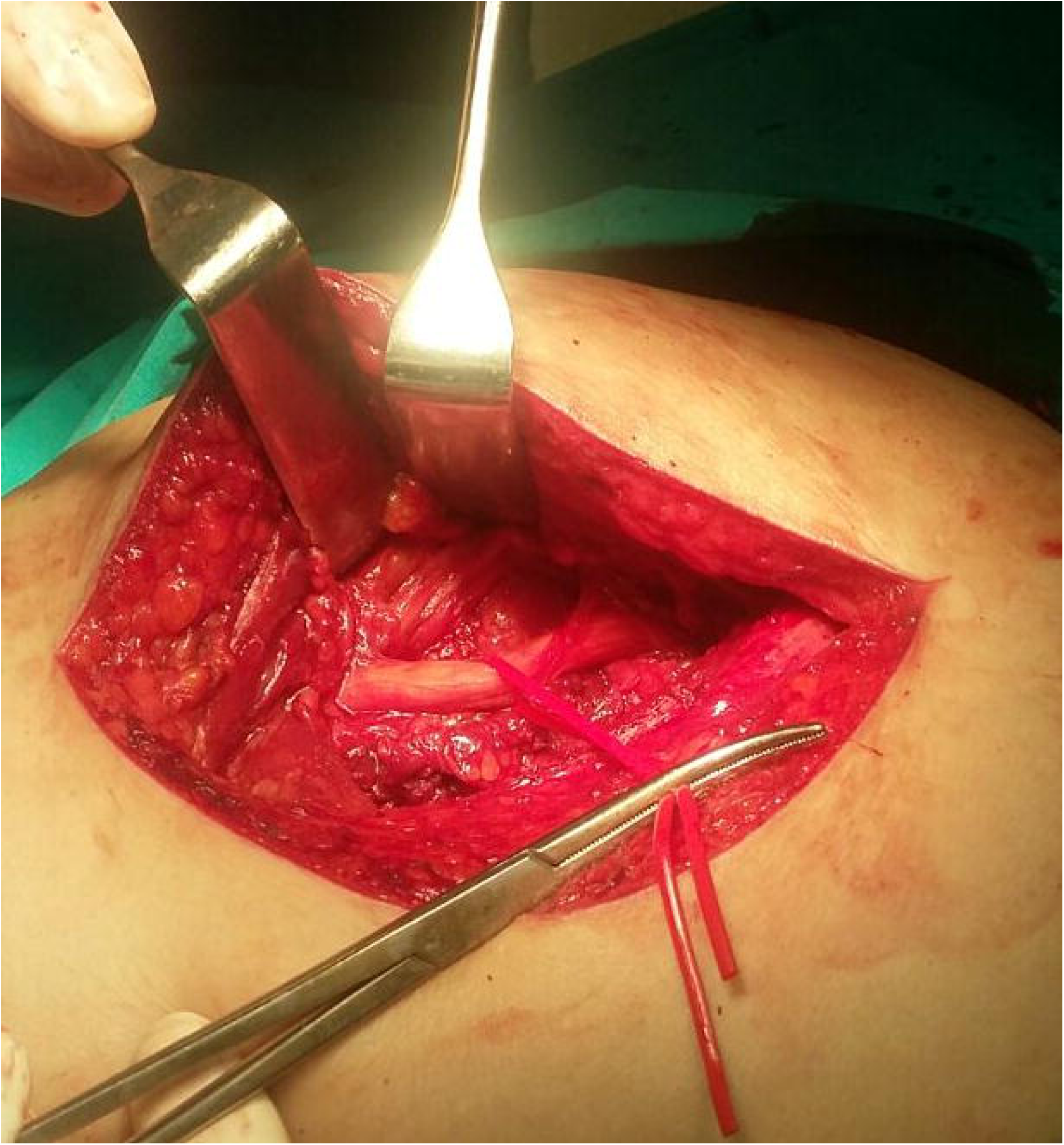
The sciatic nerve surgical release of fibrous bands.

Case 2 - Surgical release of piriformis muscle and n. peroneus beneath it was performed. This kind of surgical treatment is classified as open surgery due to posterior approach to the joint with tenotomy of m. pyriformis. Department of Orthopedic Surgery, Clinical center Zemun.

## Conclusions

The variation of the sciatic nerve is challenging for diagnostic and therapeutic procedure in many clinical and surgical cases. Quick ultrasound detection of the sciatic nerve makes surgical approaches more precise and effective, with a better outcome.

Strain elastography using ARFI seem to be a sensitive and accurate diagnostic procedure (ROC analysis) based on the assessment of nerve stiffness. This procedure could provide us with crucial information about the degree of nerve stiffness. It is a non-invasive, easily performed, reproducible and relatively inexpensive method, very useful in the process of surgical decision making and during the follow up.

